# Brain plasticity and auditory spatial adaptation in patients with unilateral hearing loss

**DOI:** 10.1101/2022.08.15.503609

**Authors:** Mariam Alzaher, Kuzma Strelnikov, Mathieu Marx, Pascal Barone

## Abstract

Unilateral hearing loss (UHL) alters binaural cues affecting speech comprehension and sound localisation. While many patients with UHL perform poorly on binaural tasks, some are able to adapt to monaural deficit. We aimed to identify patients with UHL who use compensatory strategies and to explore the neural correlates of this adaptation using Mismatch Negativity (MMN). We recruited 21 patients with UHL and we separated them into three groups using cluster analysis based on measures of binaural processing. The resulting groups were referred to as the better, moderate and poorer performers clusters (BPC, MPC and PPC). We measured the MMN elicited by deviant sounds located 10°, 20° or 100° away from a standard sound. We found that the BPC group had a significant MMN for all three deviant sounds, as in a group of normal-hearing controls. In contrast, the PPC group and normal-hearing controls with an earplug did not have a significant MMN for the 10° and 20° deviations. For the 100° deviation, the scalp distribution was found to be maximal over central regions in the BPC group, while the PPC group showed a more frontal distribution. Differences were also found for the N100 evoked by standard sounds, with the BPC group showing a contralateral pattern of activation, as in the controls, and the PPC group showing more symmetrical hemispheric activation. These results indicate that patients with UHL can develop adaptive strategies that are reflected by sound processing differences at the cortical level.

## Introduction

Binaural integration is an important function for sound localization and speech seregation. Interaural time and intensity differences (ITD and ILD) are the main cues engaged in horizontal localization (Grothe 2010), in addition to monaural spectral shape cues used predominantly for vertical and front/back localization. Accordingly, unilateral Hearing Loss (UHL), leads to an alteration of interaural balance, consequently altering spatial hearing in the horizontal plane. This alteration is expressed by higher signal to noise ratios (SNR) to understand speech in noisy environment and by a shift of the auditory space towards the better ear (Firszt et al., 2015; Rothpletz et al., 2012; Vannson et al., 2015). Middlebrooks and colleagues investigated auditory spatial adaptation after UHL (Makous & Middlebrooks, 1990) and found that patients displayed variable spatial accuracy for sound localisation, ranging from near-normal to a total shift of localisation towards the hearing side (Middlebrooks & Green, 1991). In contrast, normal-hearing controls showed poor accurate sound localisation under monaural listening conditions. Thus, UHL patients have highly variable speech reception thresholds (SRT). In a study by (Vannson et al., 2017) the SRT score in a speech-in-noise test (French Matrix test) were variable between subjects, some of them were considered as normal performers. This inter-individual variability can be attributed to individual compensatory strategies, which may differ between patients. The adaptive processes could involve a range of mechanisms, including improved processing of monaural spectral shape cues and head-shadow effects (HSE), (Batteau, 1967; Slattery & Middlebrooks, 1994; Van Wanrooij & Van Opstal, 2005) both of which contain auditory spatial cues (Agterberg et al., 2011; Van Wanrooij & Van Opstal, 2004).

The acoustic compensatory mechanisms used by patients with UHL reflect a potential for neuroplasticity (Alzaher et al., 2021). While there are evidences that human adult subjects can adapt to monaural hearing (Keating et al 2016), the brain adaptation has been only reported in animal models trough electrophysiological recording (Keating et al 2013, 2015). In humans, functional imaging studies have shown cortical processing changes in UHL patients. Specifically, while normal-hearing controls display higher activity in the hemisphere contralateral to a monaural stimulus (Elberling et al., 1981; Pantev et al., 1986), UHL patients show more symmetrical activation patterns (D. Bilecen et al., 2000; Ponton et al., 2001). However, there are currently few studies that have investigated auditory cortical plasticity for spatial hearing. In a previous study, we presented evidence that the functional integrity of the auditory dorsal stream is affected by UHL, and that this contributes to the sound localisation deficits (Vannson et al., 2020). However, further work is required to explore neural plasticity in patients.

Studies on event related potentials (ERPs) have shown that the auditory mismatch negativity (MMN) can be a useful marker of auditory dysfunction (Näätänen et al., 2015). To elicit a MMN, repetitive sound stimuli, are presented, leading to a decline in the neural response due to a passive store of repetitive information (Duncan et al., 2009). Whenever a rare change is introduced to this repetitive sequence a new ERP response is automatically elicited (Näätänen & Alho, 1997). The difference between these rare-stimulus ERPs and the repetitive-sound ERPs yields the MMN waveform. It has been found that the MMN occurs in response to a variety of stimulus disparities, including the sound spatial displacement (Deouell et al., 2006; Freigang et al., 2014).

In this study, we analysed the MMN elicited by changes in sound location (spatial MMN) to assess the functional reorganization and compensatory brain plasticity in patients with UHL. We hypothesized that MMN can be a neural correlate that reflects spatial auditory behaviour, and that MMM components would be associated with the inter-individual variability in spatial hearing skills. The approach was considered to be well-suited for gaining insight into the neural mechanisms inducing auditory plasticity in some patients with UHL.

## Materials and methods

### Participants

We recruited 20 normal-hearing subjects (NHS; 10 women), aged 20–35 years, and 21 patients with UHL (12 women), aged 25–75 years. The participants underwent audiometric testing to determine the pure tone average (PTA) thresholds for the following frequencies: 250, 500, 1000, 2000 and 4000 Hz. NHS with a PTA exceeding 15 dB were excluded from the study (n = 0). All of the NHS had normal and symmetrical levels of hearing. The patients with UHL had normal hearing levels in the healthy ear, with an average PTA of 14.7 dB HL (standard deviation: 7.9), and moderate-to-profound hearing loss in the deaf ear, with an average PTA of 82.27 dB HL (standard deviation: 29.1); the average PTA difference between the healthy and deaf ears was 67.5 dB (standard deviation: 32). Written informed consent was obtained from all of the participants before the experiment. The NHS were offered financial compensation following their participation in the study.

### Sound localisation and speech-in-noise tasks

For the sound localisation task, participants were seated in the centre of a semi-circular array (1m radius) of 12 loudspeakers, which were positioned from 82.5° on the subject’s right to - 82.5° on the subject’s left, with 15° between each speaker. The participants held a tablet with a picture of the surrounding speaker array on the screen. They were instructed to select the speaker that corresponded to the location of the sound. The acoustic stimulus was white Gaussian noise, band-pass filtered between 300 and 1200 Hz. We choose this stimulus as it correspond to the one we used in the MMN protocol (see below) based on previous MMN studies (Bennemann et al., 2013)(Freigang et al., 2014),. Before the sound localisation task, participants underwent a training session where they were instructed to listen to the sound coming from different loudspeakers at different times. The sound locations were simultaneously shown on the tablet in order to familiarize the subjects with the sound space. For the localisation test, the sound was presented through each of the 12 speakers in a random order, with a total of two sound presentations for each speaker. The subjects were asked to select the sound location on the tablet after each stimulus. The NHS performed the task under two conditions: a binaural condition and a monaural condition. For the latter, an earplug was used to produce an attenuation of 37 dB, and we added an earmuff that gives 37 dB attenuation; this resulted in a total attenuation of 40 dB.

Speech-in-noise segregation (SpiN) was evaluated by determining speech reception thresholds (SRT) using the French Matrix test (Jansen et al., 2012). This test consists of 50 different pre-selected words that fall into five categories: 10 names, 10 verbs, 10 numerals, 10 objects and 10 colours. Random combinations of one word from each category produce 5-word sentences, such as “Charlotte attrape douze ballons verts” (translation: Charlotte catches 12 green balls), which are read out by a female French speaker. In our study, the sentences were presented in variable levels of competing noise under three different conditions: (1) a diotic condition, where the speech and noise were presented through a single loudspeaker located in the centre of the array, (2) a dichotic condition, where the speech and noise were presented separately through two different loudspeakers positioned at 60° and – 60°, with the noise presented through the speaker closest to the better ear, (3) a reversed dichotic condition, identical to the dichotic condition, but with the speech presented through the speaker closest to the better ear. The participants were asked to repeat the words that they heard. If the answer was correct, the level of the noise was increased by 5 dB; if the answer was wrong, the level was reduced by 5dB. The volume of the speech remained constant throughout the test and was set at 60 dB.

### EEG recordings

Participants were seated in an armchair in a sound attenuated room, which had sound-absorbing foam to reduce echoes. EEG recordings were obtained while a series of sounds were presented through four loudspeakers. These were arranged in a semi-circular array around the participants, at a distance of 90 cm and level with the participants’ ears. An oddball paradigm was used, designed to evoke a MMN, with the standard sound presented through a loudspeaker located at 50°, and three deviant sounds, that differed only in terms of the spatial location, presented through speakers positioned at 60° (10° difference), 70° (20° difference) and –50° (100° difference; see figure 1). A total of 2000 sounds were randomly presented through the four loudspeakers, with a probability of 85% for the standard position (with a minimum of 6 repetitions before the deviant), and 5% for each of the three deviant positions.

**Figure 1.**
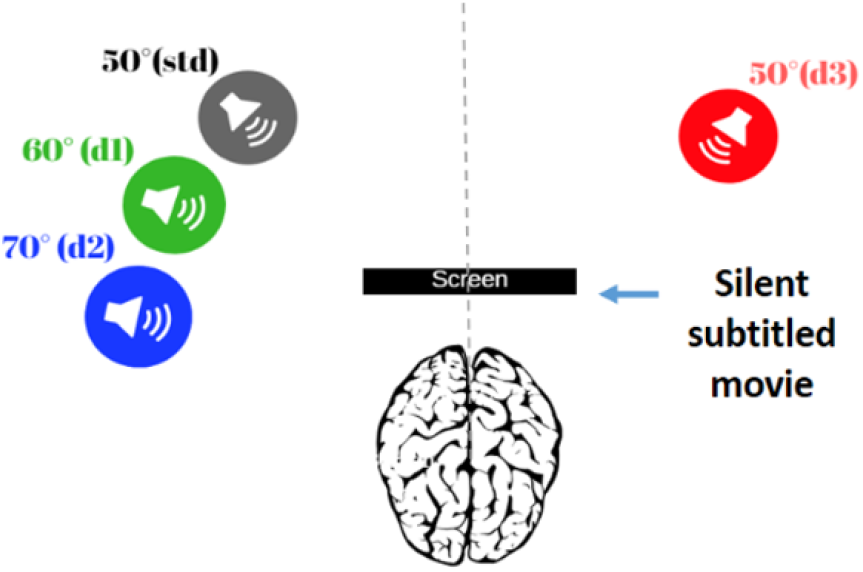
EEG auditory stimulation setup. The standard sounds were presented through a loudspeaker located at 50° (grey), and the deviant sounds (oddballs) were presented through loudspeakers located at 60° (green), 70° (blue), and -50° (red). The participants were asked to ignore the sounds and to focus on a silent, subtitled movie during the recording session.

### Properties of acoustic stimulation

The choice of stimulation type (frequency, intensity and duration) in MMN experiment is very crucial because MMN characteristics (amplitudes and latency) depend on the magnitude of difference between the physical sound properties of standard and deviant (Näätänen & Alho, 1997). Therefore, it was important to control all the physical properties to make sure of eliciting a MMN strictly related to spatial deviation. The sound duration was set at 100 ms including a 10 ms fade in and 10 ms fade out (Duncan et al., 2009). The ISI slightly varied between 490 and 500 ms to prevent stimulus onset prediction (Bennemann et al., 2013). For frequency, it was important to avoid a major intervention of intensity cues, that are more likely to elicit a MMN response related to intensity differences (Paavilainen et al., 1989), we used a low frequency band noise band-pass filtered between 300 and 1200 Hz (Bennemann et al., 2013; Freigang et al., 2014). For similar concerns, we decided to avoid an intensity roving to minimize deviances that are not strictly related to auditory spatial attributes (Cai et al., 2015; Deouell et al., 1998, 2006, 2007; Näätänen & Alho, 1997; Paavilainen et al., 1989). Therefore the level of stimulation was calibrated at sound level of 65 dB at the subjects head. During the EEG recordings, the participants watched a silent, subtitled movie on a tablet screen placed at a distance of 30 cm directly in front of them. They were instructed to ignore the sounds and to read the subtitles of the silent movie.

The 20 NHS were randomly divided into two subgroups, with 10 subjects in each group. For both of these groups, EEG recordings were obtained twice: once for a binaural condition and again for a monaural condition, where hearing was attenuated on one side using an earplug and earmuff. For the first subgroup, the hearing attenuation was on the side contralateral to the standard sound (contralat condition); for the second subgroup, the attenuation was ipsilateral to the standard sound (ipsi-condition). For the patients with UHL, EEG recordings were also obtained twice in a random order across patients: once with the deaf ear contralateral to the standard sound (contralat-condition), and again with the deaf ear ipsilateral to the standard sound (ipsi-condition). The patients’ better-hearing ear is mainly stimulated in the contralat-condition, which enables lateralisation patterns of cerebral activation to be evaluated; in the ipsilateral condition, spatial discrimination can be determined for the auditory hemifield that is most impacted by the hearing loss. Although the stimulation was not strictly monaural, due to varying degrees of residual hearing in the deaf ear, we reasoned that the stimulation at 50° would only be perceived by the better-hearing ear in the contralat-condition. This reasoning is based on: (1) the intensity of the standard sound (65 dB at the sound source, 42 dB at the subjects’ ipsilateral ear), (2) the attenuation of the bandpass noise due to the head-shadow effect (head-shadow attenuation ∼ 4 dB) and (3) the degree of hearing loss in the deaf ear (PTA ranging from 36.5 dB HL to total deafness). Therefore, the residual hearing in the deaf ear would not have affected the responses recorded at the cortical level in the contralat-condition, because the sound levels did not exceed the auditory thresholds for that ear, with the exception of one patient (who would have perceived a sound of 1.5 dB at the deaf ear).

The EEG recordings were obtained using an Active 2 system with 64 Ag/AgCl electrodes placed according to the international 10-20 system. The CMS/DRL electrodes were designated as the ground electrodes. Additional electrodes were placed at the right and left mastoids. The impedance of the scalp electrodes was kept below 10 kΩ, and the default sampling rate was used (2048 Hz).

### Data analysis

#### Analysis of the behavioural data

For the sound localisation task, the Root Mean Square (RMS) error was determined for each spatial position. The average RMS error for all 12 locations was also determined for each participant. To determine whether there was an effect of group on the average RMS error, we used a linear mixed-effects model (lme4) in R followed by post-hoc multiple comparisons using the glht function in R. The average RMS was calculated for the six speaker locations ipsilateral to the better ear (non-deaf/unplugged) and for the six locations ipsilateral to the poorer ear (deaf/plugged); these values were fitted in a seperate linear mixed-effects model (lme4) to determine whether there was an effect of side on the localisation performance (followed by post-hoc comparisons). The level of significance was set at p < 0.05.

For the SpiN results, analyses were also run using a linear mixed-effects model (lme4 in R), with group and condition as the main factors. Multiple post-hoc comparisons were then run between the groups and conditions using the glht function in R.

#### Analysis of the EEG recordings

The EEG data were analysed offline using EEGLAB (version 14.1.1.b) and ERPLAB (version 8.10), an open-source toolbox (Delorme & Makeig, 2004; Lopez-Calderon & Luck, 2014) that runs in Matlab (version 8.1.0.604 R2013a). The data were downsampled to 500 Hz and band-pass filtered at 1–20 Hz (1813-point Kaiser windowed-sinc FIR filter, Kaiser beta = 5.65, firfilt plugin version 1.5.3.) (Bennemann et al., 2013). Epochs were created for each sound stimulus, 600 ms in duration, including a 100 ms pre-stimulus baseline. Epochs with an amplitude exceeding 100 μv were rejected. The epochs for each loudspeaker position were averaged for each individual to generate ERP waveforms for the standard and three deviants. Difference waveforms were obtained for each subject by subtracting the ERP for the standard from the ERP for each of the deviants. The individual ERPs were averaged to generate grand averages for the standard and three deviants. The difference waveforms were also averaged to generate grand averages for the 10°, 20° and 100° deviations.

To determine the statistical significance of the MMN, we used a paired permutation test based on randomization (David et al., 2020). This was run using the negative peaks for the individual difference waveforms in a ±10 ms window corresponding to the latency peak of the grand average (GA) MMN at Fz electrode (Duncan et al., 2009). The analysis was run for each of the three deviant sound locations.

To determine whether the MMN amplitudes and latencies differed between the groups and deviant sound locations, we used a linear mixed-effects model (lme4). For this, the peak MMN amplitude and latency were determined for each individual at the Fz electrode, and these were fitted in the model with two factors: group (NHS-binaural, NHS-monaural, BPC and PPC; see below for definition) and sound deviation (10°, 20° and 100°). Post-hoc tests were run using the glht function for multiple comparisons.

Analyses were run to identify the electrode regions that had the largest MMN. As the MMN is known to be most prominent at the fronto-central electrodes, we restricted the analyses to the frontal and central scalp areas. We conducted a cluster comparison between the frontal (AF3, AFz and AF4) and central (C1, Cz and C2) electrodes, and a linear mixed-effects model (lme4) was used to analyse the peak MMN amplitude at each electrode as a function of the cluster (frontal vs central), both between and within the subject groups. Permutation tests were run to determine the significance of the MMN waveforms using Matlab version 2020b, and repeated measures tests (lme4) and post-hoc tests (glht) were run to compare the MMN responses using R library version 3.6.3 in R-Studio version 1.1.423.

#### K-means cluster analysis

K-means cluster analysis is an exploratory data analysis technique that aims to maximise intra-group homogeneity. Based on different measures, the k-means algorithm computes the Euclidean distance with respect to the centroid of the variables fitted in the algorithm in order to group those that are similar into a common cluster. As the 21 UHL patients in our study differed greatly in terms of their demographic and clinical characteristics (age, duration of deafness, PTA) and binaural listening skills, we conducted K-means cluster analysis to identify subgroups with similar binaural performance. Prior to the analysis, the optimal number of clusters was determined using the elbow method for the Within-cluster Sum of Squares (WSS) (D. Sharma, 2019).

The cluster analysis was run using the behavioural results obtained for the sound localisation and speech-in noise tasks, which both involved binaural processing. Five different measures were included in the analysis: (1) the RMS errors for the sound localisation task; (2,3,4) the three SRTs for the different speech-in-noise test conditions (described above); and (5) the Speech, Spatial and Qualities of Hearing Scale scores (SSQ; an auto-evaluation of speech and spatial auditory processing, run for each patient with UHL) (Vannson et al., 2015). The k-means cluster analysis was conducted using R library version 3.6.3 in R-studio version 1.1.423.

## Results

### Overall performance on the sound localisation and speech-in-noise tests

Before dividing the patients with UHL into clusters, we analysed the overall group differences on the sound localisation and speech-in-noise tests. A linear mixed-effects model was run (lme4) with Group as main factor and Subjects as random effect, followed by post-hoc comparisons between groups using glht function in R. It was found that the patients with UHL had higher RMS localisation errors compared with the NHS in the binaural condition (M = 32.1±SD=17.8 versus 15.6±4.8; p < 0.001). However, when the NHS wore an earplug (monaural condition), the RMS localisation errors significantly increased (39±10.4; p < 0.001). Although the level of hearing loss in the patients with UHL was far greater than the attenuation produced by the earplug (mean PTA in UHL group = 82.2±29.1 dB HL, versus 40 dB attenuation from the earplug), there was no significant difference between the patients and the NHS in the monaural condition (p = 0.18).

For the SpiN tests, it was found that the SRTs increased in the NHS after the introduction of an earplug in both the dichotic (−14.5±1.7 vs -2.5±4.6; p < 0.001) and reversed dichotic conditions (−15±1.6 vs -12.3±3.1; p < 0.01). The patients with UHL had higher SRTs compared with the NHS (no earplug) for all three conditions (p < 0.001). However, when the NHS wore an earplug, the SRTs did not differ significantly compared with the patients for the reversed dichotic condition, where the speech was ipsilateral to the healthy/non plugged ear (−12.3±3.1 vs - 10.3±3.7; p = 0.07).

### Cluster analysis

The K-means cluster analysis identified three clusters of patients with UHL: six better performers (Better Performers Cluster; BPC), nine moderate performers (Moderate Performers Cluster; MPC) and six poorer performers (Poor Performers Cluster; PPC). These three clusters were divided according to several dimensions, which correspond to the principal components that influence the distribution of the data points, as determined using centroids. In our analysis, dimensions 1 and 2 contributed the most to the division (dimension 1: 39.9%, dimension 2: 24.5%), as shown in figure 2, while dimensions 3, 4 and 5 were less involved (22.1%, 10.6% and 3%, respectively). To identify the behavioural test that most influenced the division into clusters, we analysed the correlations between dimensions 1 and 2 and each of the behavioural tests included in the cluster analysis. We found that the SpiN SRTs most influenced the cluster division, as there was a strong correlation between dimension 1 and the SRTs, especially for the diotic and dichotic conditions (table 2).

**Table 1.** Patient characteristics. Abbreviations: PTA = ‘pure tone average threshold’, D. side= ‘deafness side’, DS = ‘deaf side’, HS = ‘hearing side’, RMS = ‘average root mean square error for sound localisation’; M = ‘male’, F = ‘female’; R = ‘right ear’, L = ‘left ear’, SSQ= ‘Speech, Spatial and Qualities of Hearing Scale questionnaire’, total D.= ‘total deafness’, Cong = ‘congenital’, Ch. = ‘chronic’.

| N | Age | Sex | D. side | PTA-DS | PTA-HS | RMS | D. onset | Etiology | SSQ |
| --- | --- | --- | --- | --- | --- | --- | --- | --- | --- |
| 1 | 65 | F | R | 36.5 | 12 | 8.2 | 2004 | Ch. Otitis | 3.9 |
| 2 | 49 | F | R | 38 | 14.5 | 31.2 | 2017 | Unknown | 2.6 |
| 3 | 71 | M | L | 41.5 | 21 | 6.08 | 2014 | Trauma | - |
| 4 | 63 | F | R | 46 | 17.5 | 15.2 | 2018 | Unknown | - |
| 5 | 69 | M | R | 65.5 | 14.5 | 21.1 | 2015 | Unknown | 5.3 |
| 6 | 37 | M | L | 67 | 28.5 | 17.8 | 2012 | Trauma | 1 |
| 7 | 25 | M | L | 68.8 | 11 | 28.05 | 2012 | Cholesteatom | 6.1 |
| 8 | 41 | F | R | 71 | 28 | 24.4 | 2018 | Unknown | - |
| 9 | 67 | M | L | 72 | 32.5 | 60.6 | 2008 | Unknown | 5.08 |
| 10 | 69 | M | R | 72.5 | 12.5 | 26.15 | 2019 | Menier | 9.75 |
| 11 | 40 | F | L | 72.5 | 9 | 27.2 | 2015 | Unknown | 3.66 |
| 12 | 81 | F | L | 79.5 | 28.5 | 30.9 | 2018 | Sudden | 6 |
| 13 | 25 | F | R | 86 | 3.5 | 28.05 | 2006 | Unknown | 5.75 |
| 14 | 67 | F | R | 93.8 | 16.3 | 40.3 | 2016 | Unknown | 3.1 |
| 15 | 57 | F | L | 100.5 | 8.5 | 21.5 | 2017 | Ch. Otitis | 4.6 |
| 16 | 68 | F | L | total D. | 20.5 | 32.2 | Cong | Sudden | 5.41 |
| 17 | 35 | M | R | total D. | 7 | 37.8 | 2010 | Unknown | 6.37 |
| 18 | 25 | M | R | total D. | 1.5 | 62.5 | 2010 | Trauma | 6.16 |
| 19 | 44 | M | R | total D. | 6 | 44.2 | Cong | Cong | 3 |
| 20 | 57 | F | L | total D. | 12 | 79.8 | Cong | Cong | 1.6 |
| 21 | 48 | F | L | total D. | 15.5 | 30.7 | Cong | Cong | 9.46 |
| Mean(SD) |  |  |  | 82.2 (29) | 14.7 (8) | 32.1 (18) |  |  | 5 (2) |

**Table 2.** Correlation between the cluster analysis dimensions and the behavioural test scores and demographic characteristics. The analyses were restricted to the first two dimensions, as they contributed the most to the division into clusters (see Figure 2; dimensions 1 + 2 = 64.4%).

| Behavioral factor.Dimension | Corr | p-value | Demographic factor.Dimension | Corr | p-value |
| --- | --- | --- | --- | --- | --- |
| RMS.Dim1 | 0.32 | 0.14 | PTA.Dim1 | 0.28 | 0.21 |
| RMS.Dim2 | 0.4 | 0.06 | PTA.Dim2 | 0.63 | 0.002* |
| SSQ.Dim1 | 0.3 | 0.18 | Age.Dim1 | 0.4 | 0.06 |
| SSQ.Dim2 | 0.6 | 0.002* | Age.Dim2 | -0.31 | 0.16 |
| Diotic.Dim1 | 0.84 | 1.671e-06* | Deafness duration.Dim1 | -0.07 | 0.75 |
| Diotic.Dim2 | -0.39 | 0.07 | Deafness duration.Dim2 | -0.41 | 0.059 |
| Dichotic.Dim1 | 0.81 | 7.512e-06* |  |  |  |
| Dichotic.Dim2 | 0.44 | 0.04* |  |  |  |
| Rev-Dich.Dim1 | 0.65 | 0.001* |  |  |  |
| Rec-Dich.Dim2 | -0.55 | 0.009* |  |  |  |

**Figure 2.**
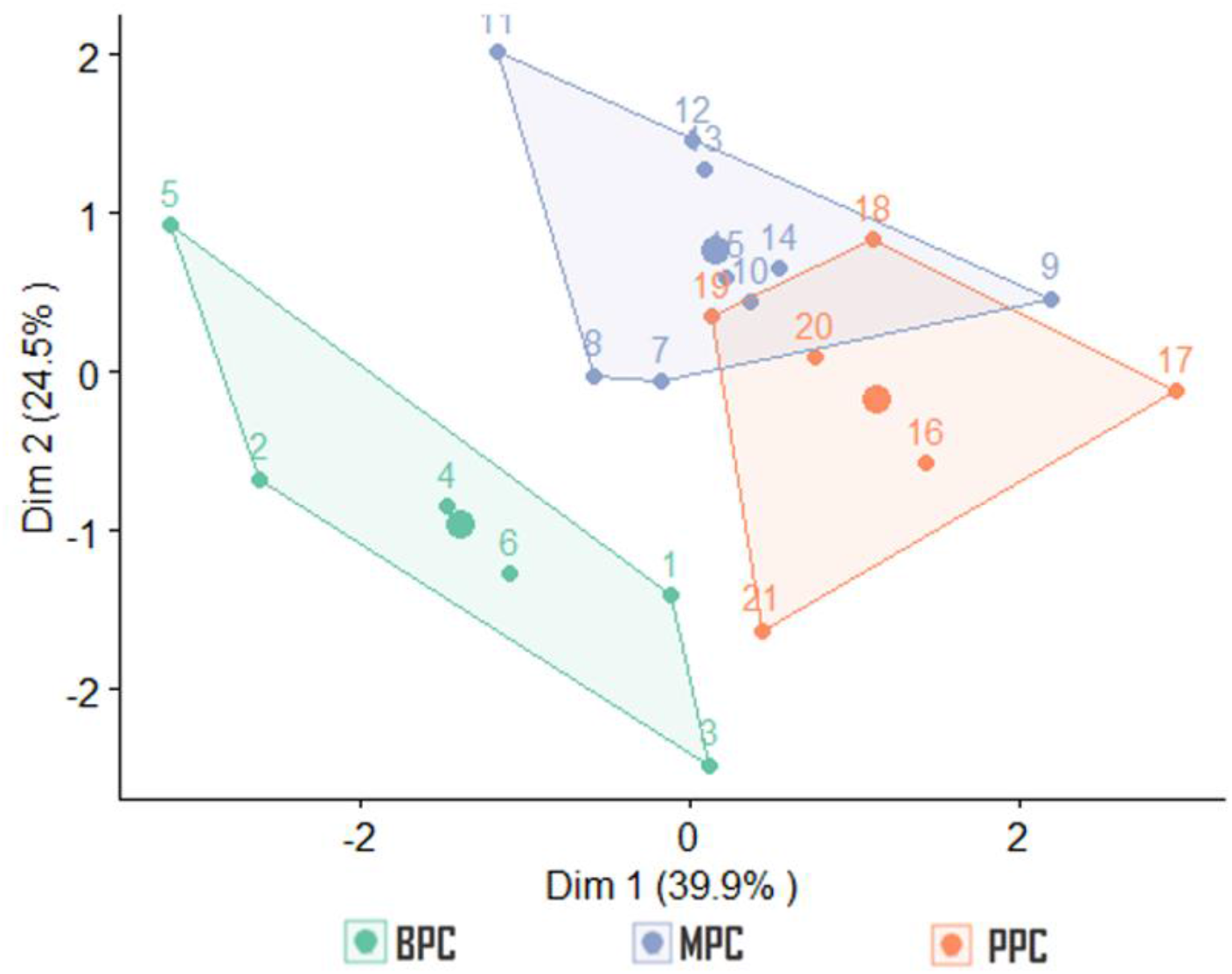
Division of 21 patients with UHL into three clusters: Better Performers Cluster (BPC), Moderate Performers Cluster (MPC) and Poor Performers Cluster (PPC). The large dots in the middle of each cluster represent the centroids. Dimension 1 had the largest influence on the division (39.9%).

To determine whether demographic factors differed between the clusters, we ran correlation analyses between dimensions 1 and 2 and certain clinical and demographic characteristics (PTA, duration of deafness and age). It was found that dimension 2 correlated with the PTA (p < 0.005), but the other correlations were not significant. To examine this further, we analysed the difference in PTA levels between the three clusters using a linear mixed-effects model (lme4) followed by post-hoc comparisons (glht function). The PTA was found to be significantly lower in the BPC group (50.8±16.3 dB HL) compared with the MPC group (97±24.1 dB HL; p < 0.005) and PPC group (91.6±22.8 dB HL; p < 0.05); the difference between the MPC and PPC groups was not significant (p = 0.88).

We ran further analyses to examine differences in the auditory test scores between the three clusters. For this, we used a linear mixed-effects model followed by post-hoc comparisons (lme4 and the glht function in R Studio). It was found that the BPC group had the lowest RMS errors (18.7±10.4) and the lowest SRT in the diotic (−3 ±3.7) and dichotic (−1.2 ±1.8) conditions compared with the MPC group (RMS: 30.4±7.4 (p=0.2) ; diotic SRT: -1.9±1.5 (p=0.6); dichotic SRT: 4.2±1.6 (p<0.001)) and the PPC group (RMS: 48.7±23 (p<0.001); diotic SRT: 1.65±2.3 (p<0.05); dichotic SRT: 5.22±2.3 (p<0.001)). For the reversed dichotic condition, there were no significant differences between the three clusters. Thus, the correlation analysis presented in table 2 shows high contribution of SRT in dichotic condition in the separation of the clusters. In these conditions, the head shadow effect (HSE) is strongly involved, therefore we suggest that BPC is the group that benefits the most from HSE compared to MPC and PPC.

### Performance on the sound localisation test for the patient clusters

We compared the sound localisation RMS for the three patient clusters (BPC, MPC and PPC) as well as the NHS group under binaural or monaural listening conditions (NHS-bin, NHS-mon). The linear mixed-effects model showed a main effect of group (p < 0.001). Post hoc comparisons showed that there was no significant difference between the NHS-bin (15.6±4.8) and the BPC (18.8±10.4) groups; however, with the earplug, the localisation errors significantly increased in the NHS (38.7±10.4; p <, 0.001), and the RMS errors were significantly higher compared with the BPC (p < 0.001) proving a strong adaptation to deafness, first in the BPC group despite the moderate hearing loss of 51 dB, second in MPC who had similar localisation errors as NHS after immediate ear plug. Of all the groups, the PPC had the highest localisation errors (48.7±23). This was significantly higher than the NHS-bin (p < 0.001), BPC (p < 0.001) and MPC (p < 0.01) groups, but not the NH-mon group (p=0.11; figure 3A).

**Figure 3.**
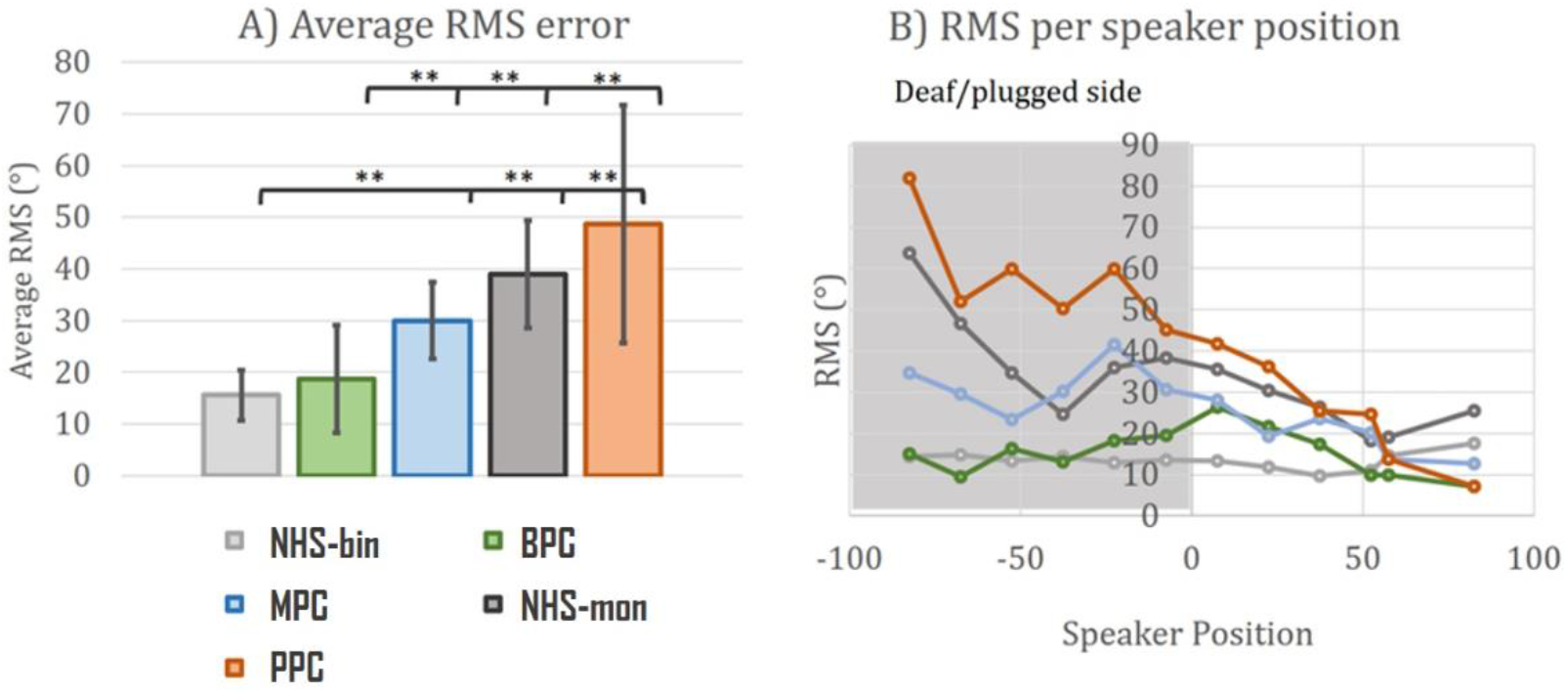
**A) RMS sound localisation errors for each group** (means and standard deviations). **B) Mean localisation errors for each group** for the different loudspeaker positions. ** p < 0.01. Abbreviations: NHS-bin= ‘ normal hearing subjects in binaural condition’, NHS-mon= ‘ normal hearing subjects in monaural condition’, BPC= ‘better performers cluster’, MPC= ‘moderate performers cluster’, PPC= ‘poor performers cluster’.

We also examined whether the side of the sound stimulation (deaf/plugged vs non-deaf/unplugged) affected the RMS localisation errors. For this, we averaged the RMS errors for the six loudspeakers on the side of the better/unplugged ear and for the six loudspeakers on the side of the plugged/deaf ear. We found that there was a significant difference between the two sides, with more localisation errors on the deaf/plugged side in the NHS-mon (40.6±13.3; p < 0.001) and the PPC (58.18±12.9; p < 0.001), but not the BPC (15.2 ± 3.6; p>0.05) compared to healthy/non plugged side (NHS-mon=25.8±6.6; PPC= 24.8±13; BPC= 15.3±7.6).

Altogether, our results are in agreement with previous data showing that UHL patient; but also NHS through training (see Florentine 1976, MacPartland et al 1997, keating 2016) can develop adaptive strategies for spatial hearing based on head shadow effect and monaural spectral cues. Based on the cluster analysis, our data emphasis that the individual patients may use different adaptive strategies which are independent of the hearing severity

### EEG results for the patient clusters

In this study, there were two monaural stimulation conditions during the EEG recordings: (1) with the standard sounds presented ipsilateral to the deaf /plugged ear, and (2) with the standard sounds presented contralateral to the deaf/plugged ear. In this section, we focus on the first of these, the ipsilateral condition, because this has a greater effect on the ability to detect spatial deviation from the standard sound position. The EEG results for the second condition, the contralateral condition, are provided in the supplementary material.

ERPs were obtained for the standard and deviant sounds and averaged for the different groups. The ERPs were characterized by a negative deflection peaking between 90 and 150 ms after the sound onset, which is known as the N100. Previous work has shown that the amplitude of the N100 varies according to the novelty of the stimulation. In the case of a repetitive stimulus (the standard), a memory trace is formed leading to a decrease in the amplitude of the N100. We were able to observe this habituation to the repetitive standard sounds in the NHS-bin and BPC groups. However, the PPC group had a higher N100 amplitude for the standard sounds, thus suggesting that each sound stimulus was processed as novel and unexpected, despite the repetition. For the NH-mon group, the N100 amplitude was small; this may relate to the binaural stimulation prior to the monaural condition, which may have led to a residual memory trace causing habituation (figure 4).

**Figure 4.**
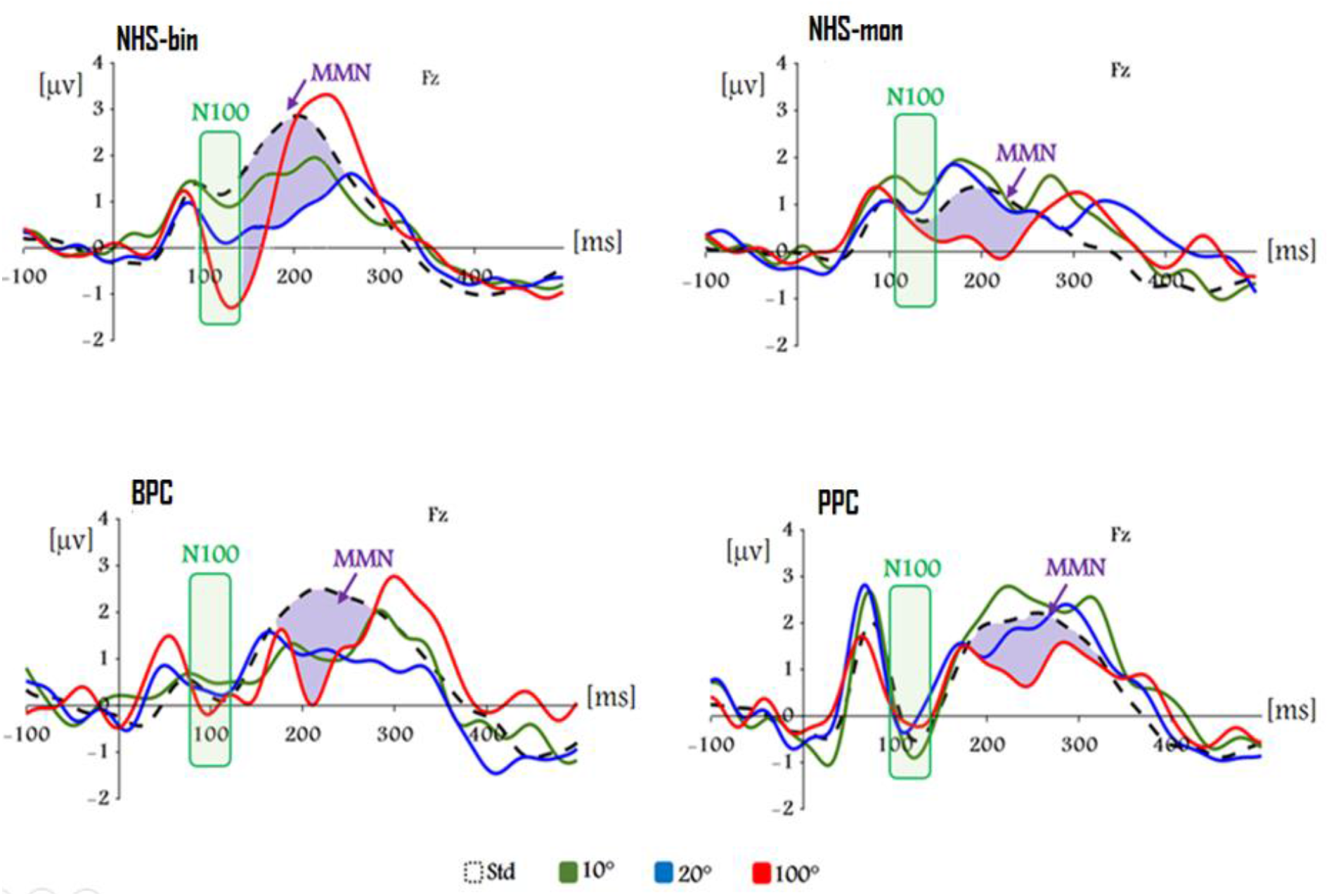
Group ERPs for standard and deviants. Grand average ERPs for the standard and deviant sounds at the Fz electrode. The separate plots are for the four groups: NHS-bin (n = 20), NHS-mon (n = 10), BPC (n = 6) and PPC (n = 6). The boxes shaded in green highlight the negative deflection of the sensory N100. The areas shaded in purple represent the MMN, which is the difference between the standard and deviant waveforms, in this case for the 100° deviation [6]. Results for the MPC group are shown in the supplementary material. Abbreviations: NHS-bin= ‘ normal hearing subjects in binaural condition’, NHS-mon= ‘ normal hearing subjects in monaural condition’, BPC= ‘better performers cluster’, PPC= ‘poor performers cluster’.

**Figure 5.**
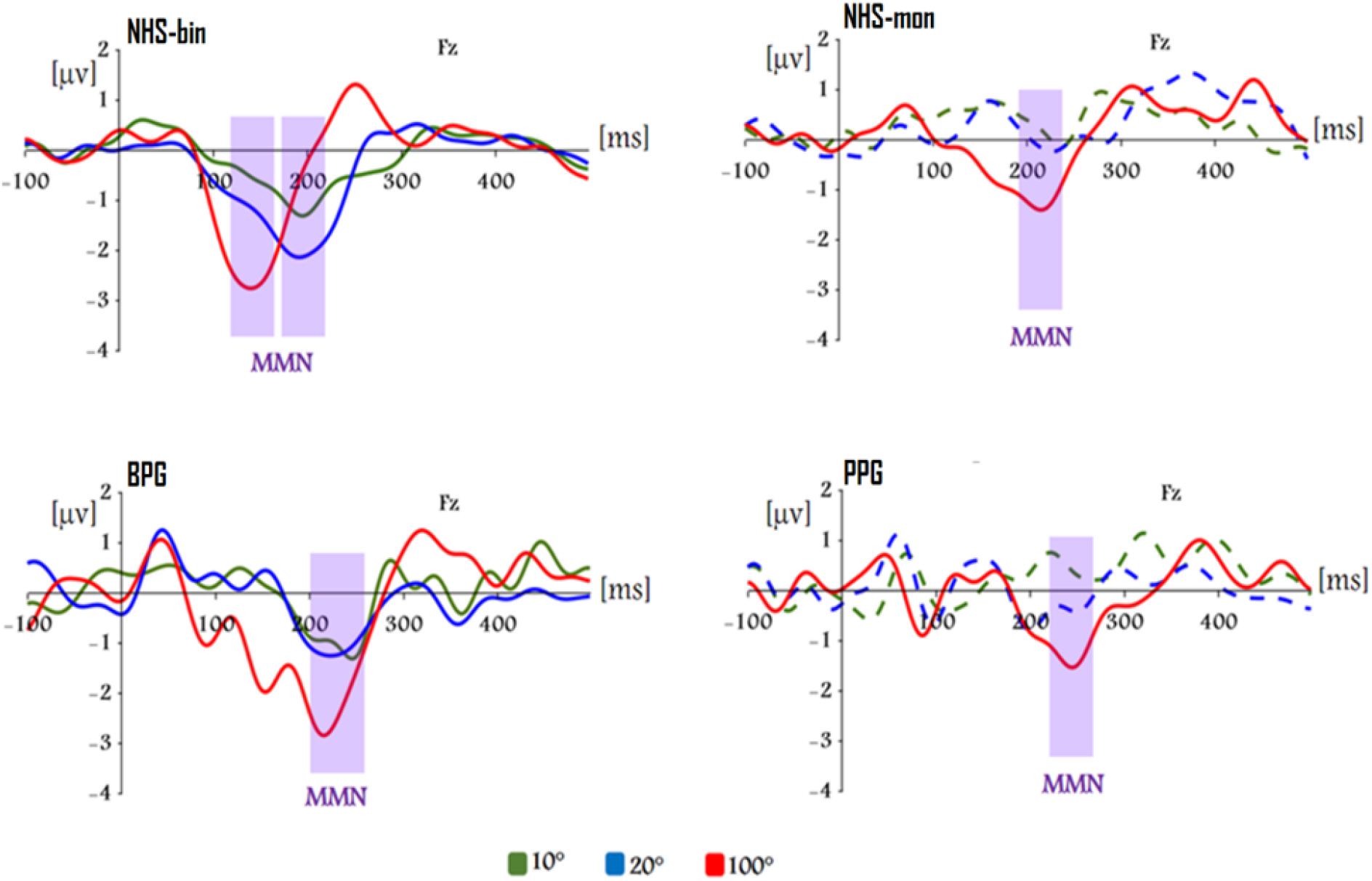
Difference waveforms for the 3 deviations. Difference waveforms (deviant ERP – standard ERP) at the Fz electrode for each of the three deviant sounds (10°, 20° and 100° deviation). The separate plots are for the four groups: NHS-bin (n = 20), NHS-mon (n = 10), BPC (n = 6), and PPC (n = 6). The dashed lines show the difference waveforms that did not have significant negativity in the permutation test. Abbreviations: NHS-bin= ‘ normal hearing subjects in binaural condition’, NHS-mon= ‘ normal hearing subjects in monaural condition’, BPC= ‘better performers cluster’, MPC= ‘moderate performers cluster’, PPC= ‘poor performers cluster’.

**Figure 6.**
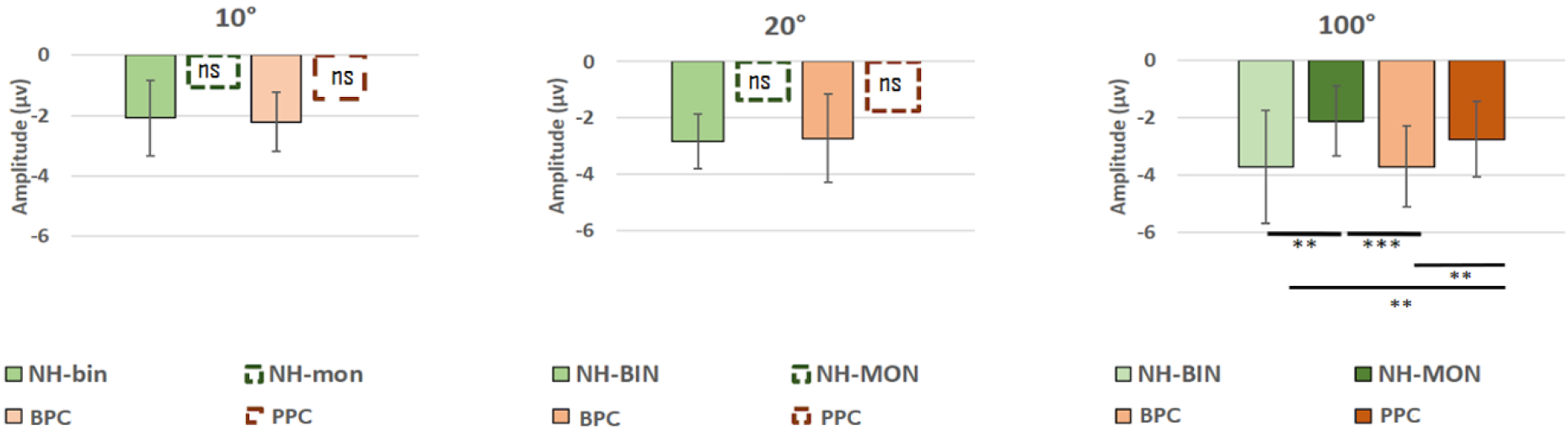
Comparison of mean individual MMN peaks at 3 deviation positions. Mean MMN amplitude peaks at the Fz electrode for the 10°, 20° and 100° deviations. The means and standard deviations are shown for each of the four groups: NH-bin (n = 20), NH-mon (n = 10), BPC (n = 6), and PPC (n = 6). If the MMN amplitudes were not significant, according to a permutation test, the means are shown using dashed lines. The asterisks show statistically significant differences based on multiple comparisons with the glht function in R. Abbreviations: NHS-bin= ‘ normal hearing subjects in binaural condition’, NHS-mon= ‘ normal hearing subjects in monaural condition’, BPC= ‘better performers cluster’, PPC= ‘poor performers cluster’, ns= ‘not significant’; ** p < 0.01; *** p < 0.001.

We observed a MMN in the ERP difference waveforms (deviant - standard), characterized by a negative deflection peaking between 150 and 250 ms. The amplitude and latency of the MMN varied according to the magnitude of the spatial deviation from the standard position. The amplitude difference between the groups and conditions was taken to reflect the spatial sensitivity of auditory processing, with higher MMN amplitudes reflecting greater sensitivity to spatial change. We therefore expected to find differences in the MMN responses for the different patient clusters, in line with their performance on the sound localisation task.

We tested the statistical significance of the MMN for the three deviant sounds in each group (NHS and the three UHL clusters). This was carried out using a permutation test based on randomization for the waveforms at the Fz electrode. Note that the MMN amplitude is determined after the N100 latency window to prevent the results from being affected by differences in the N100 for the standard and deviant sounds. It was found that a MMN was present for all three deviant sound locations in the NHS-bin and BPC groups (p < 0.05). In the MPC group, a MMN was present for the 20° and 100° deviations; in the NH-mon and PPC groups, it was only present for the 100° deviation (p < 0.05). The MMN amplitudes were compared between the different groups, but data were only included if a MMN had been found to be present. The MMN amplitudes and latencies for each group are shown in table 3.

**Table 3.** MMN amplitudes and latencies at the Fz electrode for each spatial deviation (10°, 20° and 100°) for the five groups (NH-bin: n = 20, NH-mon: n = 10, BPC: n = 6, MPC: n = 9, PPC: n = 6). Data are only shown if a significant MMN had been identified at the Fz electrode (using a permutation test based on randomisation for the individual MMN peaks in a window ±10 ms around the grand average peak). Abbreviations: NHS-bin= ‘ normal hearing subjects in binaural condition’, NHS-mon= ‘ normal hearing subjects in monaural condition’, BPC= ‘better performers cluster’, MPC= ‘moderate performers cluster’, PPC= ‘poor performers cluster’, ns= ‘not significant’, AOD= ‘angle of deviation’, p > 0.05.

| Group | AOD | Peak MMN latency<br>(Grand average) | Peak indiv. MMN amp.<br>(10 ms around latency peak) |
| --- | --- | --- | --- |
| NH bin | 10° | 194 ms | -2 μV ±1.2 |
| | 20° | 192 ms | -2.8 μV ±0.9 |
| | 100° | 140 ms | -3.7 μV ±1.9 |
| NH mon | 10° | ns | ns |
|  | 20° | ns | ns |
| | 100° | 214 ms | -1.3 μV ±1.4 |
| BPC | 10° | 246 ms | -1.5 μV ±1 |
| | 20° | 222 ms | -1.7 μV ±1.1 |
| | 100° | 216 ms | -3 μV ±1.8 |
| MPC | 10° | ns | ns |
| | 20° | 194 ms | -0.5 μV ±0.8 |
| | 100° | 200 ms | -1.1 μV ±1.2 |
| PPC | 10° | ns | ns |
|  | 20° | ns | ns |
| | 100° | 244 ms | -1.8 μV ±0.6 |

A linear mixed-effects model was run to analyse the individual MMN peak amplitudes. This showed a progressive increase in the MMN with increasing deviation from the standard sound location for the group with normal binaural hearing (Table 4). Analyses were also run for the MMN latency, using the glht function in R for multiple comparisons to study latency variation according to the angle. This revealed a decreased MMN latency for the 100° deviation (138±24.9 ms) compared with the 10° deviation (199.4±62.2 ms; p < 0.05) in the group with normal binaural hearing (n = 20); there was also a tendency for a decreased latency for the 100° deviation compared with the 20° deviation in these subjects (190.4±58,4 ms; p = 0.06).

A significant MMN was present for the 100° deviation in all of the groups. The amplitude of this MMN was higher in the NHS-bin and BPC (−3.6 μv ±1.4) groups compared with the NHS-mon (−2.1 μv ±1.2) and PPC (−2.7 μv ±1.3) groups (p < 0.05); the NHS-bin and BPC groups did not differ significantly.

We ran a further analysis to determine whether the MMN latencies differed between the five groups (NHS-bin, NHS-mon, BPC, MPC and PPC). For this, the NHS-bin group was restricted to the 10 participants who later underwent ipsilateral monaural stimulation (ipsi-condition), as we were testing for a group effect. The analysis was run using the latency of the individual MMN amplitude peaks in a linear mixed-effects model (lme4). It was found that there was no significant main effect of group (p = 0.12; figure 7).

**Figure 7.**
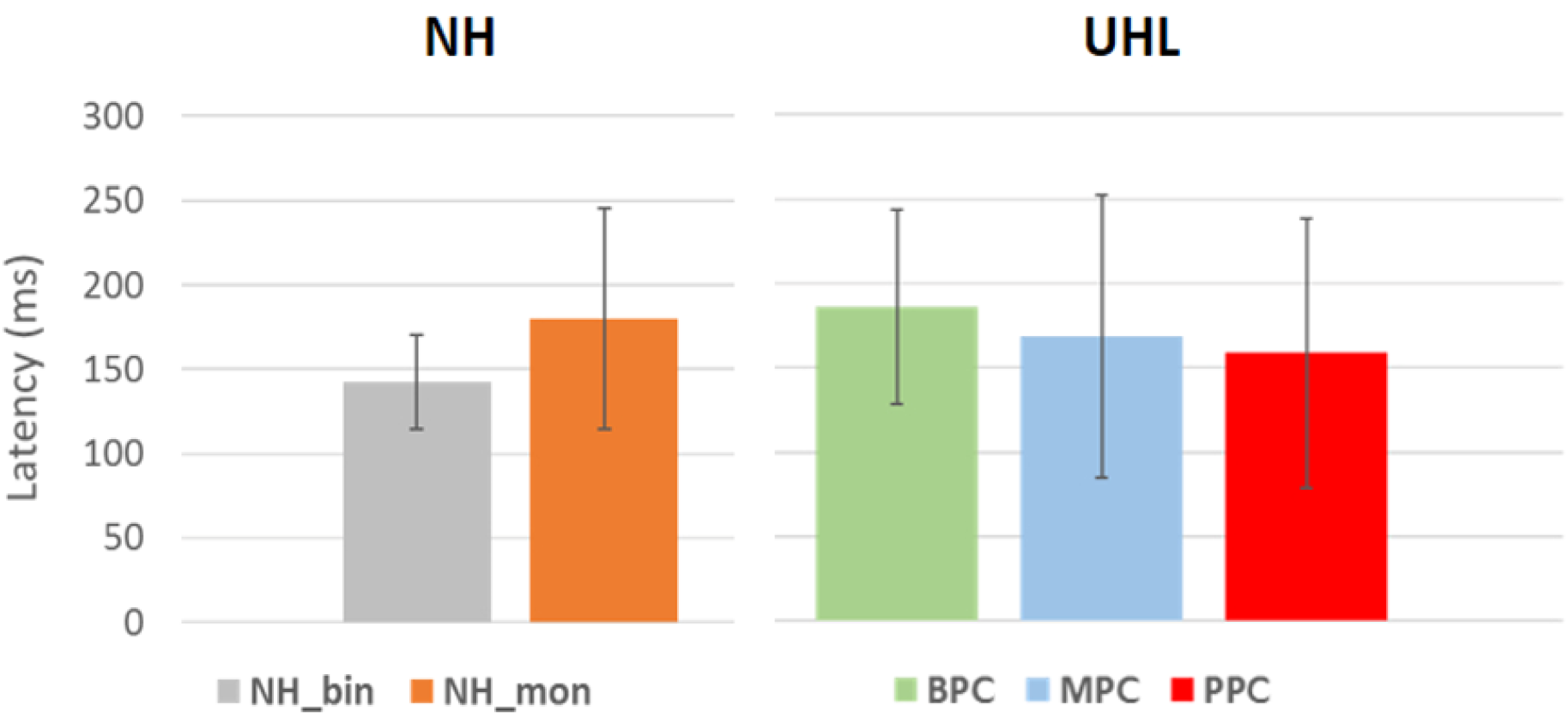
Peak latency for 100° of deviation. MMN latency at the Fz electrode for the 100° sound deviation. The latencies were determined for each participant at the peak MMN amplitude. The means and standard deviations (error bars) are shown for each group (NHS-bin, n = 10; NHS-mon, n = 10; BPC, n = 6; MPC, n = 9; PPC, n = 6). There was no significant difference between the five groups (p > 0.05). Abbreviations: NHS-bin= ‘ normal hearing subjects in binaural condition’, NHS-mon= ‘ normal hearing subjects in monaural condition’, BPC= ‘better performers cluster’, MPC=‘moderate performers cluster’, PPC= ‘poor performers cluster’, ns= ‘not significant’; ** p < 0.01; *** p < 0.001.

### Topographic distribution of the MMN

We analysed whether variation in the scalp distribution of the MMN relates to deafness or the adaptation to deafness. Previous work has shown that the MMN is maximal at the frontal and central electrodes (Fz and Cz) (Näätänen et al., 2017; Zhang et al., 2011). Higher activation at the central electrodes has been related to attentional switching due to the automatic detection of deviants (Opitz, Rinne, Mecklinger, Yves Von Cramon, et al., 2002), while lower activation has been related to a limited ability to discriminate or integrate surrounding sounds. Higher activation at the frontal compared to the central areas has been related to the cognitive demands involved in ignoring auditory stimuli and concentrating on a set task (Campbell & Sharma, 2013). For our analyses, we therefore focused on three frontal electrodes (AF3, AFz and AF4) and three central electrodes (C1, Cz and C2). We restricted our analysis to the 100° sound deviation because the MMN was significant for all five groups and the large deviation would have led to more marked attentional switching.

We ran a linear mixed-effects model with cluster and range of interest (frontal vs central) as factors, and subjects as random effect, followed by a post-hoc comparisons (glht function in R). The scalp distribution of the MMN is shown in figure 8 for the NHS-bin, NHS-mon, BPC, MPC and PPC groups. The patterns of activation were observed to be similar for the NHS-bin and BPC groups, characterized by higher central electrode activation and lower frontal electrode involvement.

**Figure 8.**
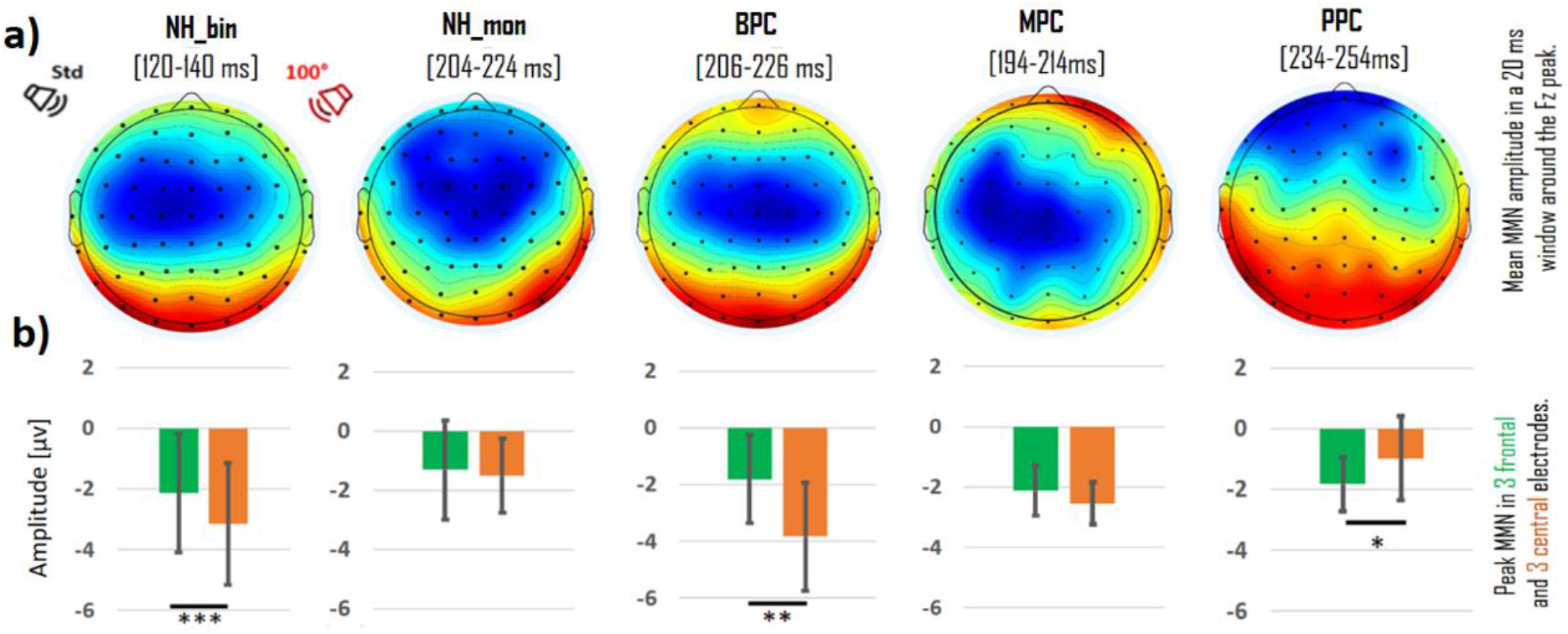
Scalp distribution of the MMN evoked by a 100° deviation in sound location. The plots are based on the mean MMN amplitude in a 40 ms time window centred on the MMN peak. The bar charts show the mean amplitudes of the individual MMN peaks at the three frontal (AF3, AFz, AF4) and three central electrodes (C1,Cz,C2). Abbreviations: NHS-bin= ‘ normal hearing subjects in binaural condition’, NHS-mon= ‘ normal hearing subjects in monaural condition’, BPC= ‘better performers cluster’, MPC=‘Moderate performers cluster’, PPC= ‘poor performers cluster’;*p<0.05; ** p < 0.01; *** p < 0.001.

The NHS-mon group showed a different pattern, with equal activation at the frontal and central regions (−1.3±1.6 vs -1.5±1.2; p = 0.52). The PPC group presented a reversed pattern of activation compared with the NHS-bin and BPC groups, characterized by weak activation at the central electrodes and higher activation at the pre-frontal areas (−1.8±0.8 vs -0.9±1.3; p < 0.05).

Thus, the central electrode activations were higher in the NHS-bin (−3.1±1.9) and BPC (−3.8±1.9) groups compared with the NH-mon (−1.5±1.2; p < 0.01) and PPC (−0.9±1.3; p < 0.01) groups.

For the MPC we observed a pattern between that seen for the BPC group (greater activity over the central regions) and the PPC group (greater activity over the frontal regions). There was no significant difference between the MMN amplitudes at the frontal and central regions in the MPC group, unlike the PPC and BPC groups, a pattern that was also seen in the NHS with an earplug.

### Interhemispheric asymmetry of the N100

To further explore differences in the cortical activation patterns for the different UHL clusters, we analysed responses to the standard sound in the contralateral condition (when the unplugged/better ear was located on the same side as the standard sound, which was positioned at 50°). For this, we focused on the distribution of the N100 across the scalp and its lateralisation.

Figure 9 shows the scalp distribution of the N100 in a 20 ms time window centred on the N100 peak at the Fz electrode. It can be seen that the BPC and NHS-mon groups showed higher contralateral activation as a response to the standard sound, but this was not apparent for the MPC and PPC groups.

**Figure 9.**
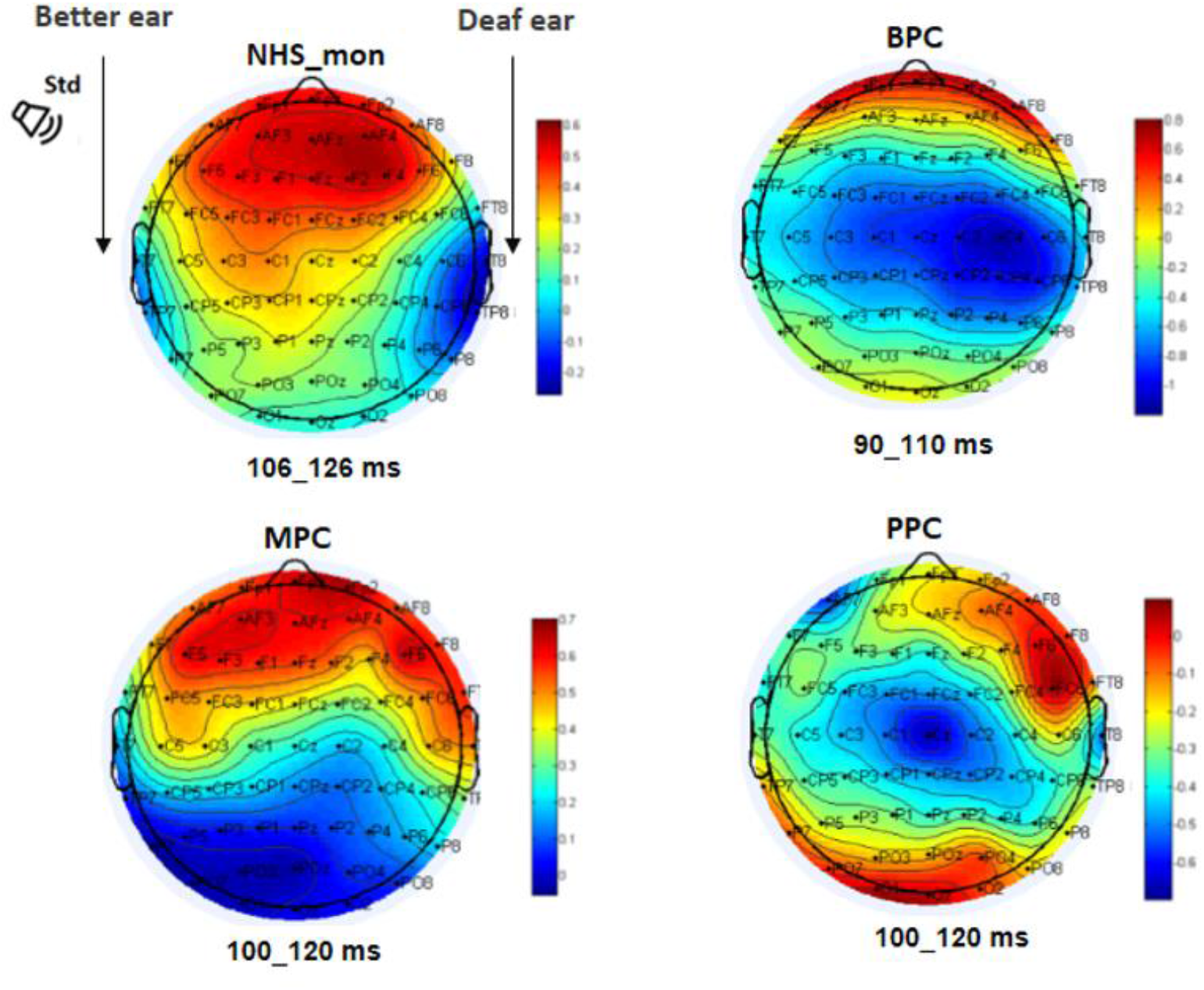
Scalp distribution of the N100 response. The results are shown for a 20 ms time window centred on the N100 peak, as determined at the Fz electrode. This peak is positive for the NHS-mon and MPC groups (see supplementary material Figure 13).

To analyse these hemispheric asymmetry patterns, we calculated an Asymmetry Index (AI). For this, nine pairs of electrodes were identified over the N100 generators, referred to as the Region Of Interest (ROI), as shown in figure 10b. Asymmetry indices were calculated for each of these electrode pairs, using the individual N100 peak values, with the following formula: AI = (contralat response – ipsi response / contralat response + ipsi response)

**Figure 10.**
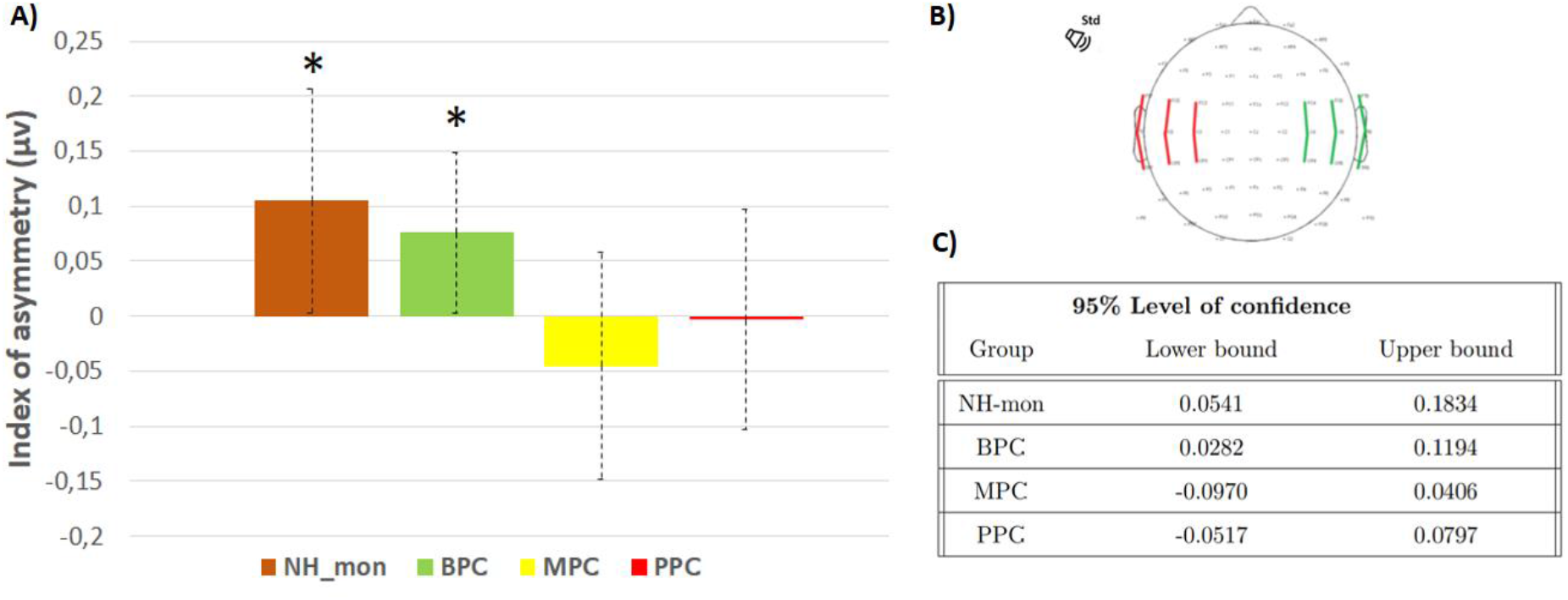
Comparison of asymmetry indexes between groups. a) Asymmetry index values for the NH-mon, BPC, MPC and PPC groups. These were calculated based on the N100 peak values at nine electrode pairs located over the N100 generators. The means and standard deviations are shown. B) Region of Interest (ROI) for the asymmetry index. C) Confidence intervals for the individual asymmetry indices, determined using bootstrap analysis. Abbreviations: NHS-bin= ‘normal hearing subjects in binaural condition’, NHS-mon= ‘normal hearing subjects in monaural condition’, BPC= ‘better performers cluster’, MPC=‘moderate performers’, ‘PPC= ‘poor performers cluster’; ** p < 0.01; *** p < 0.001.

**Figure 11.**
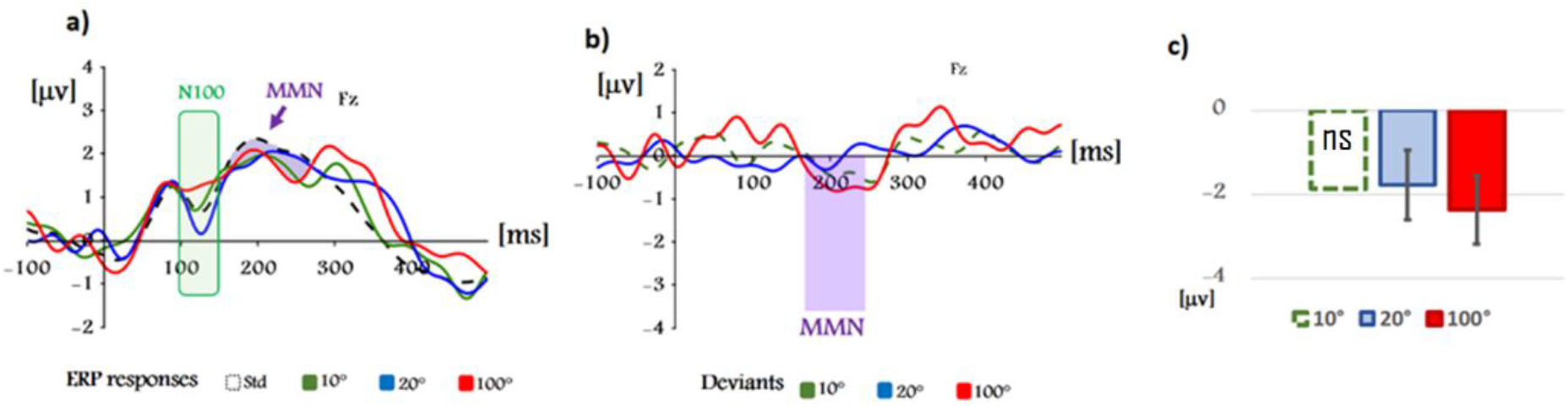
ERP and MMN responses in MPC. a) Event-related potentials for the standard and deviant sounds in the MPC group (n = 9). Results are shown for the Fz electrode in the ipsilateral condition (deaf side ipsilateral to the standard sound). B) Difference waveforms (deviant – standard) for the three different deviant sounds. The dashed line for the 10° deviation shows that the MMN was not significant according to a permutation test (p < 0.05). c) MMN peaks at the Fz electrode (means and standard deviations for the individual values). The dashed line shows that the MMN was not significant (ns).

**Figure 13.**
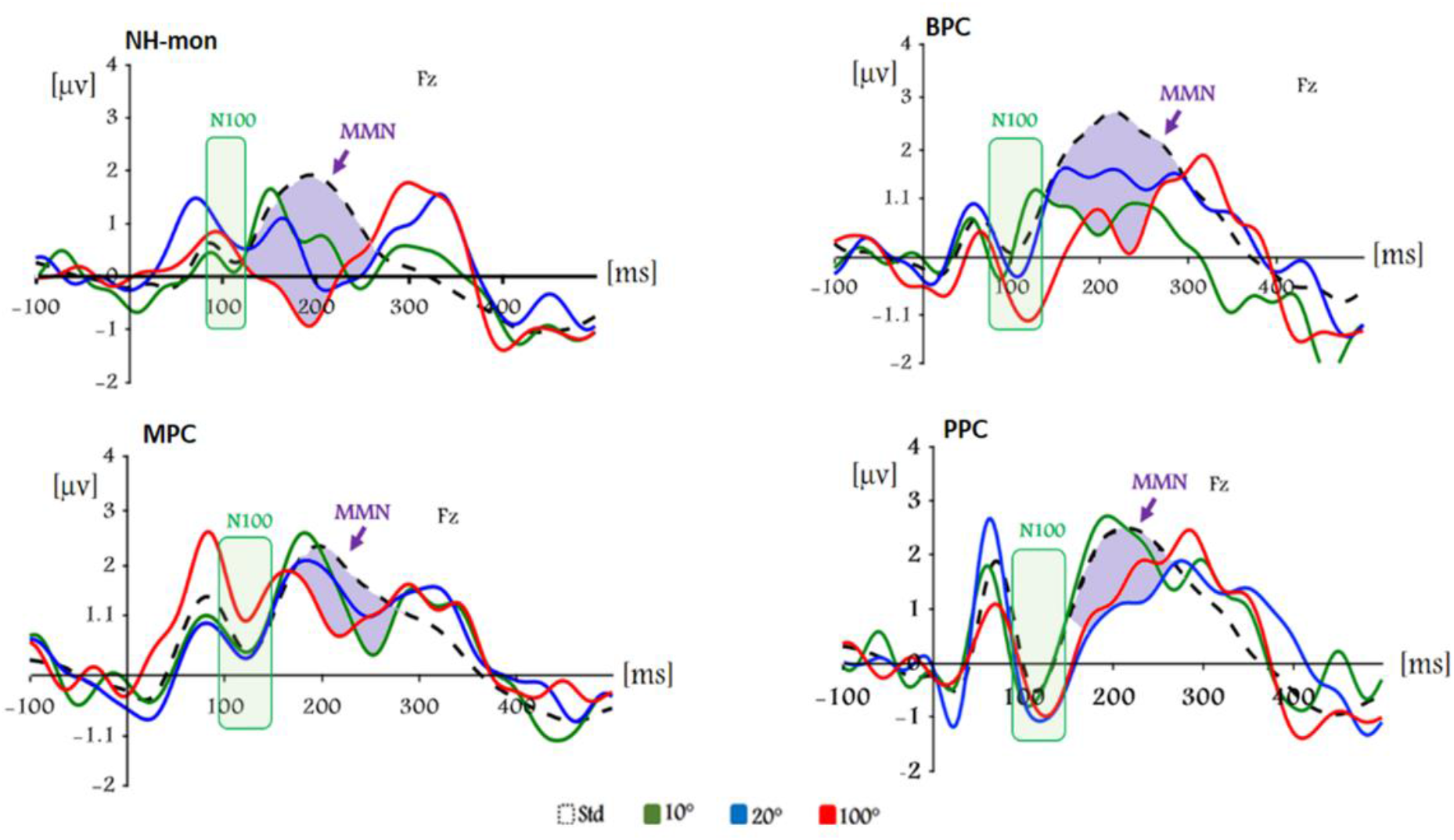
Group ERPs for standard and deviants in contralateral condition. Data are shown for the Fz electrode in four groups: NH-mon (n = 10), BPC (n = 6), MPC (n = 9) and PPC (n = 6). The boxes shaded in green highlight the negative deflection of the N100. The areas shaded in purple represent the MMN, which is the difference between the standard and deviant waveforms [6]. Note that the MMN amplitude is determined after the N100 latency window to prevent the results from being affected by differences in the N100 for the standard and deviant sounds.

**Figure 14.**
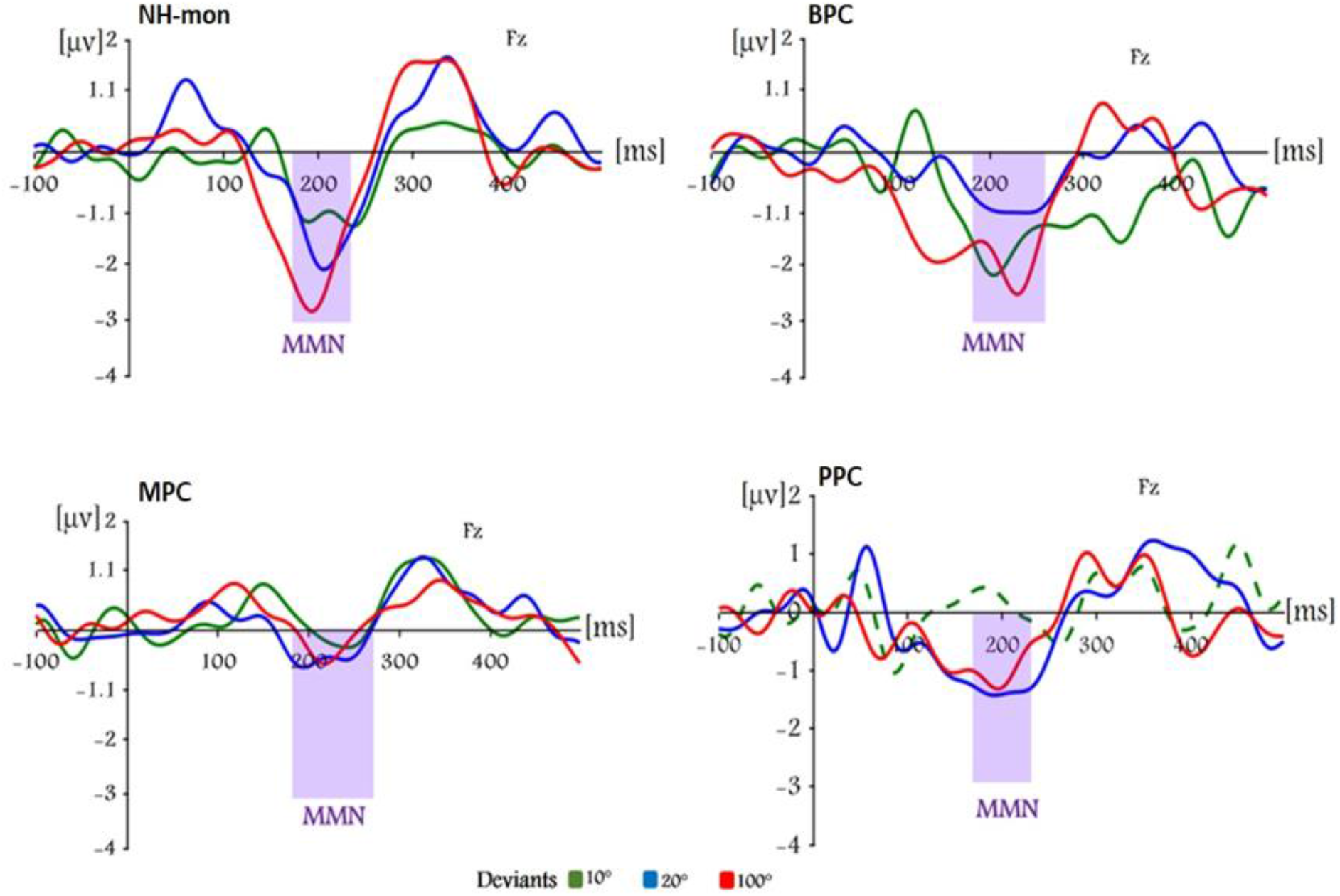
Difference waveforms at the Fz electrode for the contralateral condition. The separate plots are for the four groups: NHS-mon (n = 10), BPC (n = 6), MPC (n = 9) and PPC (n = 6). A permutation test based on randomisation was run using the individual MMN peaks (identified in a ±10 ms window around the grand average peak). This revealed a significant MMN for the three deviant positions for all groups except the PPC group, which did not have a significant MMN for the 10° deviation (green dashed line).

**Figure 15.**
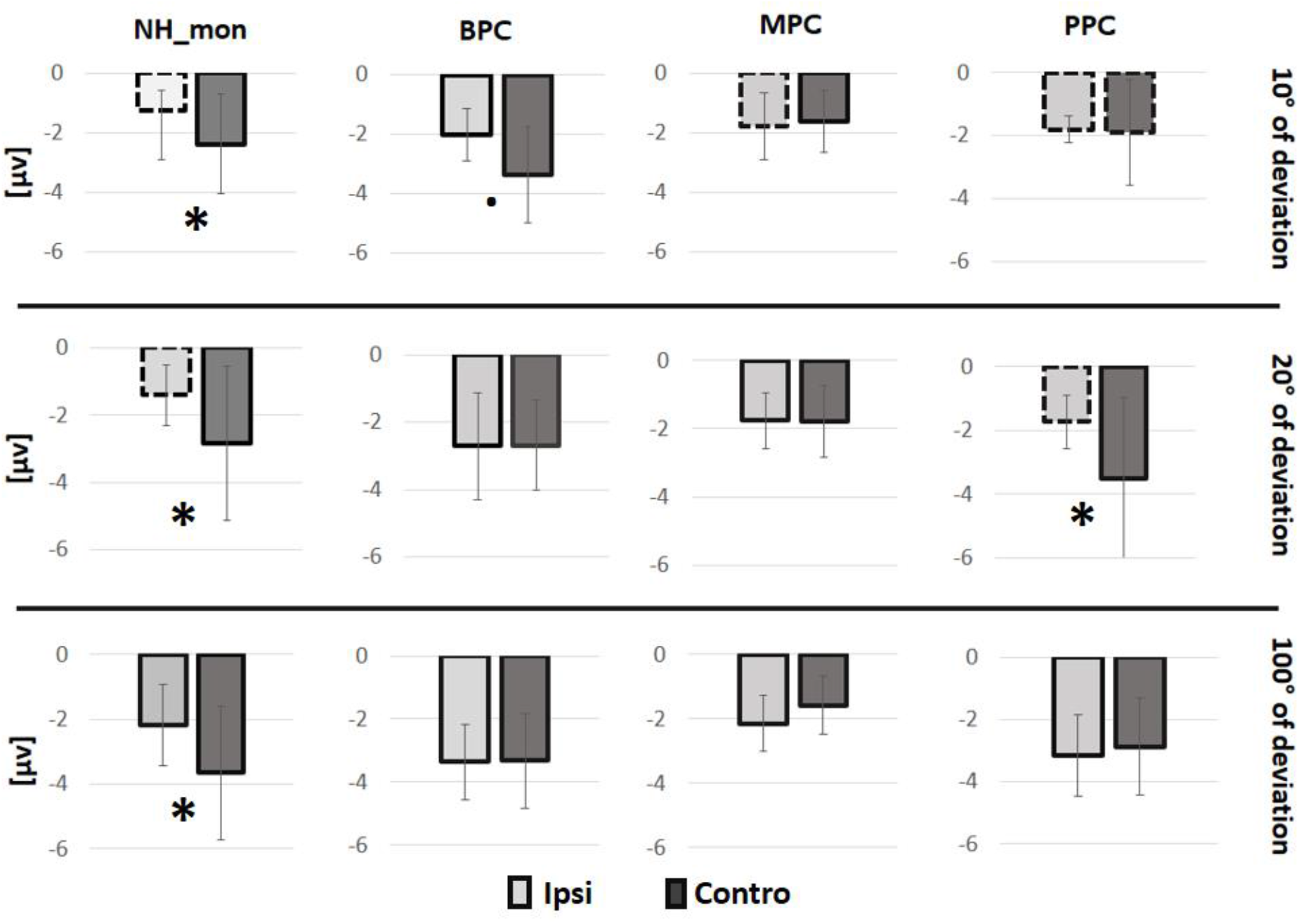
Average individual MMN peak amplitudes at the Fz electrode for the three deviant sounds. Results are shown for the four groups (NHS-mon, BPC, MPC and PPC) for the ipsilateral (ipsi) and contralateral (contralat) conditions. Dashed lines represent a non-significant MMN according to a permutation test based on randomization. Significant differences between the contralateral and ipsilateral conditions are shown by an asterisk (* p <0.05).

When the index is positive, this indicates that there is an asymmetrical pattern of contralateral cortical activation. The overall AI was calculated by averaging the AIs for all nine pairs of electrodes in the ROI.

We found that the overall AI values were positive for the NHS-mon and BPC groups, whereas they were negative or close to zero for the MPC and PPC groups (figure 10a). A bootstrap analysis revealed that the AI values were significantly positive (different from zero) for the NH-mon and BPC groups, but not for the MPC and PPC groups. These results indicate that auditory stimuli predominantly activate the contralateral hemisphere in the NH-mon and BPC groups. This is expected based on the functional lateralisation of the auditory system. In contrast, the MPC and PPC groups had a more symmetrical activation pattern, which is characteristic of the central reorganization observed in patients with unilateral deafness.

## Discussion

Studies on deafness have classically focused on how the degree of hearing loss affects auditory processing. Many studies have explored how profound deafness affects the auditory processing pathway and its effects on speech comprehension, spatial discrimination and sound localisation. However, relatively few studies have considered the neural processes that follow partial hearing loss. Here, we investigated the neural reorganisation in moderate-to-severe UHL and how this relates to different levels of auditory performance. We adopted a new approach to identify subgroups of patients with similar binaural auditory skills, which involved clustering analysis revealing a subset of patients with near-normal auditory spatial abilities and a near-normal MMN, in spite of their significant hearing loss (PTA = 51 dB HL; BPC). These patients display the classic contralateral pattern of auditory processing in response to sounds at the better-hearing ear. At the other end of the spectrum, we identified a subgroup of patients with marked auditory spatial deficits (PPC; PTA = 91 dB HL), a lack of the usual contralateral pattern of auditory activation while the MMN exhibited an accentuated frontal distribution, which may relate to cognitive deficits induced by the deafness.

### Spatial hearing and unilateral hearing loss

The patients in our study had unilateral hearing loss ranging from moderate (PTA ∼40dB HL) to profound (>90dB HL, including total deafness), and they varied greatly in terms of their spatial hearing deficit. Although hearing loss level (PTA) was not fitted in the clustering model, level of hearing loss was found to correlate with the spatial deficit as reported (Firszt et al., 2013; Heinrich et al., 2015; Humes et al., 1980; Nelson et al., 2019; Vannson et al., 2017). However, some patients with severe hearing loss (PTA > 70dB HL) were found to have normal levels of performance on sound localisation (RMS) or speech-in-noise (SRT) tasks (e.g., patients 8, 9 and 15; table 1), while others with total unilateral deafness had higher performance levels than monaural controls. These results are in line with our previous study (Vannson et al., 2017), where we identified two subgroups of patients who differed in terms of their SpiN performance with no significant difference in their PTA, suggesting that PTA cannot be the only criterion to explore auditory adaptation.

In the present study, a cluster analysis was run using the performance levels on tasks involving binaural processing. This meant that the separation of patients into subgroups did not depend on the degree of hearing loss in a strictly linear fashion. Indeed, the patients in the MPC and PPC groups had similar levels of hearing loss (severe-to-profound; mean levels: 97 and 92 dB HL, respectively), although the patients in the BPC group had a lower mean PTA (moderate-to-severe; mean level: 51 dB HL). Despite the similar hearing levels, the MPC and PPC patients differed in terms of their performance on the SpiN and sound localisation tasks; the BPC patients were found to perform as well as NHS.

### Brain plasticity and adaptation in unilateral deafness

We linked binaural performances to neural events evoked by changes in the sound location. In hearing impaired patients, the ERPs evoked by different acoustic features are translated on the behavioural level by similar levels of discrimination for those features (Cai et al., 2015; Ponton & Don, 1995; A. Sharma et al., 1993). Similar links were also found in our study : the BPC group had a mean RMS error of 15° and a MMN sensitive to 10° of deviation; in the PPC group and monaural controls, the average RMS was around 50° and the MMN was correspondingly absent for sound separations of 10° and 20°. In contrast, a MMN was present for all groups in the case of a 100° deviation. All together, we demonstrate that MMN is an accurate neural marker of spatial hearing skills.

Auditory processing is characterised by a contralateral ear dominance, i.e. representation of the contralateral sound field in each hemisphere (Phillips & Gates, 1982). In UHL patients, this cortical asymmetry is altered, with a dominance shift from contralateral to ipsilateral with respect to the better-hearing ear(D. Bilecen et al., 2000; Deniz Bilecen et al., 1998). We previously demonstrated (Vannson et al., 2020) that the ipsilateral shift relates to sound localisation deficits, thus suggesting that the change disrupts representations of the sound field. In the present study, the cluster analysis enabled the separation of patients into different groups according to their binaural processing skills, irrespective of the degree of hearing loss. We found that the cluster with near-normal spatial skills (BPC) presented a N100 that was larger over contralateral cortex following stimulation of the better-hearing ear, as in controls. In contrast, the groups with poor skills (MPC and PPC) did not show this lateralised pattern, showing instead a bilateral response to monaural stimulation. These results support our previous findings of an ipsilateral shift following UHL (Karoui et al., 2022; Vannson et al., 2020), and are in line with our hypothesis that contralateral dominance is a functional prerequisite for accurate spatial hearing and corresponds to representations of the contralateral sound field. In further support of this, patients with UHL who received a cochlear implant for their deaf ear have been found to have both restored auditory spatial skills (Bernstein et al., 2016; Grieco-Calub & Litovsky, 2010; Vermeire & Van De Heyning, 2009; Zeitler et al., 2015) and restored contralateral dominance (Debener et al., 2008; Legris Id et al., 2018; Sandmann et al., 2015). The hearing controls with an earplug performed as poorly on the spatial tasks as the PPC, and they did not have a significant MMN for the smaller spatial deviants. However, they were found to have the normal contralateral pattern of N100 responses. Previous work has shown that UHL patients with a cochlear nerve resection display a shift from contralateral to ipsilateral dominance, but this can take several months (D. Bilecen et al., 2000; Burton et al., 2012). This can therefore account for the normal contralateral pattern seen in the controls with an earplug, as the monaural stimulation was of short duration.

It could be argued that the patients in the BPC group had undergone an ipsilateral shift in auditory processing, but that this was reversed over time through adaptation and the use of monaural spatial cues. However, this could not explain the lack of adaptation in the other patient clusters, who had similar durations of deafness. Further studies are required to understand the factors that may lead to or predict the auditory adaptation seen in certain patients.

### Potential indication of cognitive deficits in unilateral deafness

The patients in the PPC group had a marked spatial hearing deficit, which was reflected by the absence of a MMN at 10° and 20° of deviations but present at 100° deviation. Thus, MMN was largely generated in the frontal lobes in these patients (Deouell et al., 1998; MH et al., 1990). Nonetheless, the NHS and BPC groups showed higher MMN signal in the central regions. We hypothesize that this difference may relate to cognitive deficits in the PPC caused by a high demand of executive resource implication to compensate for the hearing loss.

MMN alteration is known to be associated with cognitive decline and various neurological disorders (e.g. Alzheimer’s disease and schizophrenia), this alteration is usually reflected by prolonged latencies, decreased amplitudes and/or a changed scalp topography. The MMN alteration could be linked to an impairment in involuntary attention switching or to short-term memory deficits which is necessary for the MMN to be elicited (Bennemann et al., 2013; Näätänen et al., 2004). In our study, the stronger MMN signal at the frontal scalp locations in the PPC group may relate to a deficit in short term memory processing as reported in adults and children with hearing loss (Kral et al., 2016; Lin et al., 2013), leading to a lack of passive spatial encoding in the oddball paradigm (Lin et al., 2013; M et al., 2012).

The increased MMN signal over the frontal regions seen in the PPC group, and to a lesser extent in the MPC group, may reflect an extended brain network that compensates for the auditory spatial deficit. The frontal regions have been implicated in listening effort in numerous neuroimaging studies (Opitz et al., 2002; Sörqvist et al., 2016), and deficits in speech-in-noise comprehension have been linked to stronger frontal activation in hearing-impaired patients (Campbell & Sharma, 2013).

Cognitive deficit originated from a higher cognitive load and listening effort in UHL patients (Opitz et al., 2002) leads to isolation and lack of communication when hearing deficit is untreated. Auditory rehabilitation could have a beneficial role in delaying cognitive deficit especially in the type of patients corresponding to the MPC group who are still preserving executive functions reflected by pre attentive processing of spatial deviance. These patients could potentially benefit froma sound amplification up to 45–60 dB HL, which could improve their adaptation. Thus, audio-visual training has been shown to effectively improve spatial hearing skills and adaptation strategies (Strelnikov et al., 2011; Valzolgher et al., 2022)

## Conclusion

This study shows that a substantial proportion of patients with UHL perform at a near-normal level on auditory spatial tasks, which can be attributed to adaptive strategies to compensatr the disruption to binaural cues. The severity of the hearing loss cannot fully account for either the level of impairment on sound localisation tasks or the degree of recovery. We demonstrate that the adaptation to UHL is reflected at the cortical level, as shown by differences in the auditory spatial MMN that may relate to cognitive and memory deficits, including short term memory. Although the mechanisms that underlie the inter-individual variability remain unclear, our results imply that rehabilitation strategies would be effective either through conventional hearing aids or through perceptual learning.

## Supplementary material

### 1 Results for the MPC group (behavioural and electrophysiological)

The cluster analysis identified three subgroups of patients with UHL based on their performance on the binaural tasks. The MPC group was found to have severe-to-profound hearing loss that did not differ significantly from the PPC group (97±24 dB HL vs 92±23 dB HL, respectively). However, the MPC group had intermediate levels of performance on the sound localisation task, with significantly better scores than the PPC group (30.4±7.4 vs 48.7±23; p < 0.005) and poorer scores than the BPC group (30.4±7.4 vs 18±10.4; p < 0.005); the MPC group’s performance was similar to the NHS with an earplug (attenuation = 39 dB; 30.4±7.4 vs 39±10.4; p = 0.26)). For the SpiN task, the MPC group had better thresholds than the PPC group (diotic condition: -1.9±1.5 vs 1.6±2.3; p < 0.05) but poorer thresholds than the BPC group (−3±3.7; p < 0.05). These results show that performance on the different binaural tasks improves from the PPC group up to the BPC group.

In line with the behavioural results, the MPC group showed an intermediate MMN response to the spatial deviant sounds compared with the BPC and PPC groups. In the ipsilateral condition (standard sound ipsilateral to the deaf ear), a MMN was evoked for all three deviant sound locations in the BPC group (10°, 20° and 100°; p < 0.05), for two sound locations in the MPC group (20° and 100°; p < 0.05) and for only one sound location in the PPC group (100°; p < 0.05).

Altogether, the MPC group can be seen to have intermediate results for the behavioural (sound localisation and SpiN) and cortical (MMN) measures, which lie between the BPC/NHS-bin and PPC/NHS-mon groups. As the patients in the MPC group have severe hearing loss (97 dB HL), they have presumably developed adaptive strategies that enable improved but non-optimal spatial hearing.

### 2 MMN in the contralateral condition

Accurately locating sound sources is more difficult when sounds are presented ipsilateral to the deaf/plugged ear. The MMN was analysed for this condition (ipsilateral condition) to study the neural correlates of adaptive plasticity. However, the contralateral condition (standard sounds presented contralateral to the deaf/plugged ear) is also of interest, as this can provide further data concerning binaural integration and reveal differences between the ipsilateral and contralateral conditions.

For the ipsilateral condition, the permutation tests (see materials and methods) showed that there was no significant MMN for the 10° and 20° deviations in the NHS-mon and PPC groups; there was also no significant MMN for the 10° deviation in the MPC group. In contrast, for the contralateral condition, a significant MMN was found to be present for all deviant positions in all of the groups, with the sole exception of the 10° deviation in the PPC group. This result implies that the PPC group still had difficulty detecting small (10°) spatial deviance when the sound was presented ipsilateral to the better-hearing ear. This is not the case for the MPC group. We compared the MMN amplitudes obtained for the three deviant sounds (10°, 20° and 100°), the two conditions (ipsilateral and contralateral), and the different groups (NHS-mon, BPC, MPC and PPC). For this, a linear mixed-effects model (lme4) was run followed by multiple comparisons (glht function in R). As predicted, the side of the stimulation (deaf ear/earplug ipsilateral or contralateral to the standard sound) had a large effect for the NHS-mon group, with different MMN amplitudes for all three deviation positions (10°: ns vs -2.3±1.6, 20°: ns vs -2.8±2.2; 100°: -2.1±1.2 vs -3.6±2, for the ipsilateral and contralateral conditions, respectively). The PPC group also showed an amplitude difference between the ipsilateral and contralateral conditions for the 20° deviance (ns vs –3.5±2.5). However, the side of the stimulation did not have a significant effect in the BPC group (p > 0.05). In addition, the MMN amplitude for the 100° deviation did not differ significantly between the two stimulation sides for any of the three patient groups (p > 0.05). This finding could be attributed to head-shadow effects, which would enable the patients to detect spatial deviations that cross the midline, irrespective of the stimulation side or level of hearing loss.

## Acknowledgments

We would like to thank all our participants for taking part in the study. We also acknowledge the technical advices of Julien Tardieu for the experimental setups, and lastly we like to thank Jessica Foxton for her help providing critical reviews and corrections for the paper.

